# Ectopic expression of a maize gene is induced by Composite Insertions generated through Alternative Transposition

**DOI:** 10.1101/2020.08.10.245175

**Authors:** Weijia Su, Tao Zuo, Thomas Peterson

## Abstract

Transposable elements (TEs) are DNA sequences that can mobilize and proliferate throughout eukaryotic genomes. Previous studies have shown that in plant genomes, TEs can influence gene expression in various ways such as inserting in introns or exons to alter transcript structure and content, and providing novel promoters and regulatory elements to generate new regulatory patterns. Furthermore, TEs can also regulate gene expression at the epigenetic level by modifying chromatin structure, changing DNA methylation status and generating small RNAs. In this study, we demonstrated that *Ac/fAc* transposable elements are able to induce ectopic gene expression by duplicating and shuffling enhancer elements. *Ac/fAc* elements belong to the *hAT* family of Class II TEs. They can undergo standard transposition events, which involve the two termini of a single transposon, or alternative transposition events which involve the termini of two different, nearby elements. Our previous studies have shown that alternative transposition can generate various genome rearrangements such as deletions, duplications, inversions, translocations and Composite Insertions (CIs). We identified over 50 independent cases of CIs generated by *Ac/fAc* alternative transposition and analyzed 10 of them in detail. We show that these CIs induced ectopic expression of the maize *pericarp color 2 (p2)* gene, which encodes a Myb-related protein. All the CIs analyzed contain sequences including a transcriptional enhancer derived from the nearby *p1* gene, suggesting that the CI-induced activation of *p2* is effected by mobilization of the *p1* enhancer. This is further supported by analysis of a mutant in which the CI is excised and *p2* expression is lost. These results show that alternative transposition events are not only able to induce genome rearrangements, but also generate Composite Insertions that can control gene expression.

**Summary:** When Barbara McClintock originally identified and characterized Transposable Elements (TEs) in maize, she termed them “Controlling Elements” due to their effects on gene expression. Here we show that maize *Ac/Ds* TEs can acquire a genomic enhancer and generate Composite Insertions (CIs) that activate expression of a nearby gene. CIs are structurally variable elements that include TE termini enclosing sequences from an original donor locus, and are formed when the termini of two nearby TEs transpose during S phase from a replicated to unreplicated site. In this way, TEs may acquire genomic enhancers to generate Controlling Elements as described by McClintock.

## Introduction

Transposable Elements (TEs) are DNA sequences that can move their positions and proliferate themselves in the genomes. Wicker et al., published a unified classification system for transposable elements in 2007 (Wicker et al., 2007). There are two types of transposable elements: Class I TEs, are also called RNA elements, since their transpositions rely on RNA as intermediates; Class II TEs do not need RNA for their transpositions, therefore, they are also called DNA elements. Class II TEs can undergo standard transpositions: TE-encoded transposase binds to the termini of a single TE and facilitate the excision and insertion of the TE. In contrast, at least some Class II TEs can also undergo alternative transpositions, which involve the termini of two TEs. This mechanism was observed in various species and mediated by different TE families (Gray *et al*., 2000), including IS*10*/Tn*10* in bacteria (Chalmers and Kleckner, 1996), Tam*3* in snapdragon (Martin and Lister, 1989), *P* elements in *Drosophila* (Gray et al., 1996) and *Ac*/*Ds* elements in maize (Weil and Wessler, 1993). In this study, we focused on characterizing the products of a specific type of alternative transposition reaction driven by maize *Ac/Ds* elements. *Ac/Ds* was the first TE system discovered by Barbara McClintock in the 1940s (McClintock, 1948; McClintock, 1950). *Ac* is the autonomous element which encodes the transposase enzyme, and *Ds* is the non-autonomous counterpart that requires *Ac* transposase for transposition. Previous work in maize has shown that *Ac/Ds* can undergo two major types of alternative transpositions: Reversed Ends Transposition (RET) involves the reversely-oriented termini of two different elements on the same chromosome, while Sister Chromatids Transposition (SCT) targets the termini of two TEs located on different sister chromatids (Peterson and Zhang, 2013). Previous studies have shown that SCT can generate deletions, inverted duplications (Zhang and Peterson, 1999; 2005), sister chromatid fusions and chromosome breaks (Yu et al., 2010); while RET can generate deletions (Zhang and Peterson, 2005), direct duplications (Zhang *et al*., 2013), inversions, and translocations (Zhang and Peterson, 1999, 2004; Zhang et al., 2006). In addition, both SCT and RET can generate novel compound structures termed Composite Insertions (CIs)(Zhang *et al*., 2014; Wang et al. 2020).

In addition to generating genome rearrangements, TEs can affect gene expression in many different ways (Hirsch and Springer, 2016). For example, TE insertion in introns can alter splicing patterns, leading to new transcripts and protein products (Luehrsen and Walbot, 1990). Many studies have shown that TEs can provide novel promoters to drive expression of adjacent genes (Butelli et al., 2012). In certain conditions, TEs may provide enhancer sequences that trigger stress-induced gene expression (Makarevitch et al., 2015). Additionally, TEs may exert epigenetic effects on nearby genes, such as inducing the spread of DNA methylation from TEs to flanking sequences, thereby suppressing expression of neighboring genes (Hollister, J.D. and Gaut, B.S., 2009). Moreover, TEs may alter chromatin states and thereby influence gene expression:Eichten et al. (2012) reported increased heterochromatin and reduced gene expression in the vicinity of TE insertions.

Enhancers are important cis-regulatory elements in eukaryotic genomes. Enhancers are typically short, 50-1500 bp, and bound by transcription factors to activate gene expression (Blackwood et al., 1998). They can be located upstream or downstream of the target genes, and they may function over long distances by forming chromatin loops (Krivega and Dean, 2012). In maize, only a small number of enhancers have been identified and characterized (Oka et al, 2017). For example, the enhancer of the maize *booster1 (b1)* gene consists of multiple tandem 853 bp repeats located around 100 kb upstream of the *b1* coding sequence (Stam et al., 2002). The enhancer of *teosinte branched 1* (*tb1*), a maize domestication gene (Doebley et al., 1995), is located ∼ 60 kb upstream of the *tb1* target gene (Doebley et al., 2006). The gene *pericarp color1* (*p1*) controls biosynthesis of a red phlobaphene pigment in multiple maize organs such as pericarp, cob and silk. *p1* expression is regulated by dual enhancer sequences that are repeated at sites upstream and downstream of the *p1* coding sequence (Sidorenko et al.,1999). In this study we show that *p1* enhancers can be mobilized by alternative transposition events to activate ectopic expression of a second maize gene. These results demonstrate the potential impacts of TIR transposable elements and alternative transposition events on maize genome evolution.

## Methods

### Maize genetic stocks and screen

The progenitor allele *p1-wwB54* has a loss of *p1* function due to the deletion of the first two exons of *p1*, therefore, it has white pericarp and white cob. To screen for new RET events resulting in *p2* expression, plants of genotype *p1-wwB54* heterozygous with a *p1* null allele (*p1-ww [4Co63]*) were grown in an isolation field and allowed to pollinate with *p1-ww[4Co63]* pollen parents. The resulting ears were screened, and kernels with red pericarp were selected and propagated; heritable cases were further analyzed for insertions in *p2*. Genomic DNA samples were screened by PCR using primers located in the *Ac* and *fAc*, paired with primers from the *p2* gene sequence. Samples giving positive results for both *5’Ac/p2* and *3’fAc/p2* junctions were considered to be candidate CI alleles. These candidate CI alleles were then planted and self-pollinated to generate homozygotes for analysis. To screen for further mutations of CI *S7* and *E3* alleles, plants carrying these alleles were self-pollinated or crossed with *p1-ww [4Co63]* in the isolation field; in resulting ears, kernels with white pericarp (*S7M*) or light red pericarp (*E3M*) were selected as mutants derived from the respective CIs.

### DNA extraction and Polymerase Chain Reaction (PCR)

Total genomic DNA was prepared by using a modified CTAB (cetyltrimethylammonium bromide) extraction protocol from leaves of 3-week old plants. Promega GoTaq® Green Master Mix was used for PCR reactions. The PCR is initiated by a 2-minute denaturation at 95 °C, then 30 seconds of annealing step at the temperature of 5 °C below the melting temperature of the primers, then 1 min extension per kb at 72 °C; these steps are repeated for 30 cycles and a final extension at 72 °C for 5 minutes was applied.

### RT-PCR

Total RNA was extracted by Invitrogen TRIzol™ Reagent from maize pericarp 20 days after pollination, and treated with NEB DNase I to remove genomic DNA. cDNA was prepared by Invitrogen™ SuperScript™ II Reverse Transcriptase kit and used as the template of RT-PCR.

### Bisulfite Sequencing

Bisulfite treatment was performed using EZ DNA Methylation-Lightning Kit from Zymo Research. The bisulfite-converted DNA was used as template for PCR using primers that were designed based on the converted sequences. Sequence conversion was done by the program MethPrimer2.0 (http://www.urogene.org/methprimer2/tester-invitation.html) (Li and Dahiya, 2002).

### Data and Reagent Availability

Maize genetic stocks are available by request to TP. Sequences reported here are available in the Supplemental Materials.

## Results

### Composite insertions produced from B54 via RET during DNA replication

The *pericarp color 1* (*p1*) gene encodes a R2R3 Myb transcriptional factor, and regulates phlobaphene biosynthesis in maize floral organs including kernel pericarp and cob glumes (Dooner, H.K., 1991). *pericarp color 2* (*p2)* is a paralog of *p1*, but is not expressed in pericarp and cob. Both *p1* and *p2* are located on the short arm of maize chromosome 1, separated by ∼70 kb (Zhang et al. 2000). Phenotypes of *p1* alleles are commonly identified by a 2-letter suffix which indicates the color of pericarp and cob, respectively. For example, *p1-ww* indicates white pericarp and white cob, and *p1-rw* indicates red pericarp and white cob (Brink and Styles, 1966). The *P1-rr11* allele conditions red pigmentation of kernel pericarp and cob. It contains an intact *p1* gene with a full length (4565 bps) *Ac* element inserted upstream of *p1* exon 1, and a fractured *Ac* (*fAc*, only 2039 bp of 3’ of *Ac*), inserted in *p1* intron 2 (Figure 1) (Zhang and Peterson, 2004). In a previous study, Yu et al. (2011) showed that the *Ac* and *fAc* termini in *P1-rr11* could undergo RET to induce deletions of the DNA between the *Ac/fAc* termini. In one case, deletion of *p1* exons 1 and 2 produced a mutant allele termed *p1-wwB54* (hereafter referred to as *B54)*, with colorless pericarp and cob. The *B54* allele retains the *Ac* and *fAc* elements in reversed orientation, with the 5’ terminus of *Ac* and 3’ terminus of *fAc* separated by a segment of 331 bp (Figure 1). In this configuration, the *Ac* and *fAc* termini in *B54* can generate sister chromatid fusions and chromosome breaks (Yu et al., 2011).

**Figure 1.**
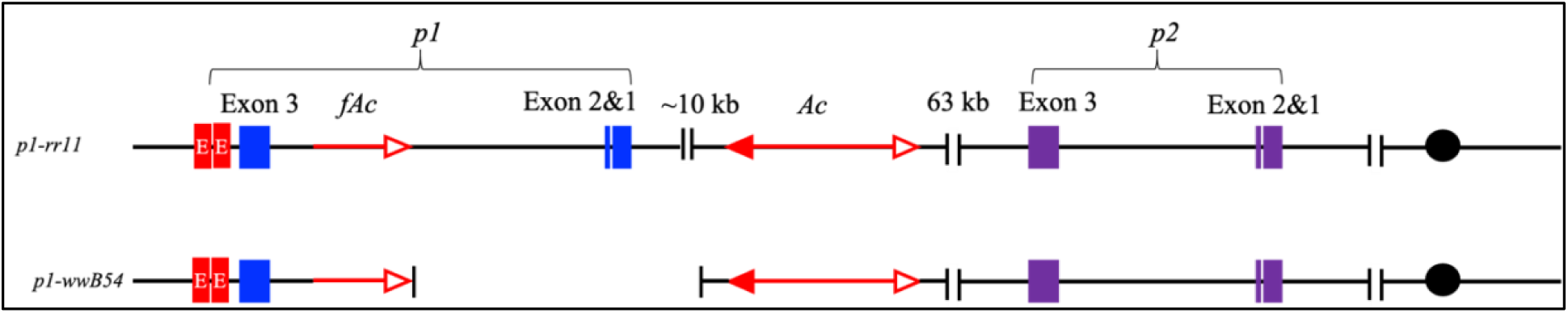
Structures of *p1-rr11* and *p1-wwB54*. The upper line indicates the structure of the progenitor allele *p1-rr11*. The lower line indicates the structure of *B54* which has a deletion of exon 1 and exon 2 of *p1*. The blue and purple boxes indicate the exons of *p1* and *p2* genes, respectively; red boxes indicate copies of enhancer *f15*. The arrow with a single open arrowhead indicates fractured *Ac (fAc);* double-headed arrow indicates full-length *Ac* element. Black dots indicate the centromere of chromosome 1.

Using a different *p1* allele, a previous study showed that a pair of reverse-oriented *Ac/fAc* in *p1* can undergo RET and induce DNA re-replication to generate flanking duplications and novel structures termed Composite Insertions (CIs) (Zhang et al, 2013; Zhang et al, 2014). (Because the formation of duplications was previously described in detail, here we focus on the formation and function of the CIs). We hypothesized that *B54* may also produce CIs via RET during DNA replication, as shown in Figure 2. In this model, the *Ac* transposase excises the 3’ end of *fAc* and 5’ end of *Ac* from a region of replicated DNA, and inserts these termini into an unreplicated target site. This insertion generates a rolling circle replicon to re-replicate *Ac* and flanking sequences, while *fAc* and its flanking sequence will be re-replicated by elongation of the impinging replicon. At some point, re-replication spontaneously aborts to produce two broken ends with Double Strand Breaks (DSBs). The fusion of these two DSBs will rejoin the two chromosome fragments and generate a CI at the new junction. If the re-replication fork through *fAc* is sufficiently extended, the CI is expected to include a copy of *p1* exon 3 and transcriptional enhancer element *f15*.

**Figure 2.**
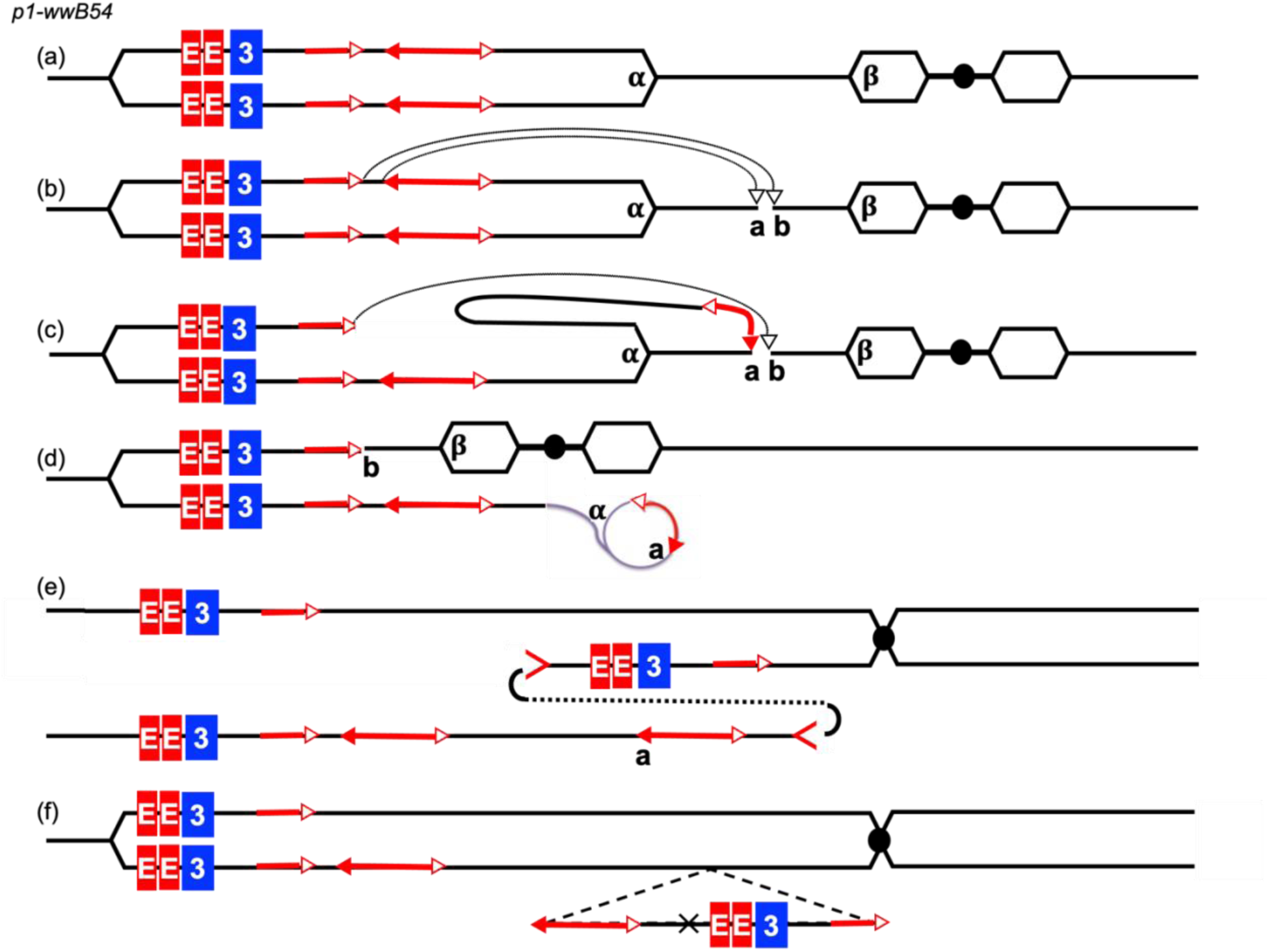
Model of CI formation from *B54*. (a) The structure of the allele *p1-wwB54*. The hexagons indicate replicons. α and β indicate two replication forks. Other symbols have the same meaning as in Figure 1. (b) Transposase binds to the *fAc* and *Ac* and the two termini insert into the target site *a/b*, which is not yet replicated. (c) The insertion of *Ac* generates a rolling circle replicon and the insertion of *fAc* joins with target site b. (d) *Ac* and its flanking sequence are re-replicated by the rolling circle replicon. (e) The re-replication aborts and the two double strand breaks (indicated by > and <) fuse together. (f) A CI is generated containing *Ac, fAc, p1* exon 3 and enhancer *f15*, and a portion of the flanking sequences.

### Unexpected reversion of deletion allele *p1-wwB54*

Initial observations of maize ears produced by plants containing *p1-wwB54* showed that many kernels contained red sectors resembling the red revertant sectors typical of somatic reversion of *p1-vv* to *P1-rr* (Emerson, 1929). This was surprising, considering that both exons 1 and 2 of the *p1* gene were deleted in *p1-wwB54*. These two exons contain most of the coding sequence for the Myb DNA binding domain that is essential for *p1* function (Grotewold et al.,1991). We hypothesized that these sectors may result from ectopic expression of the *p2* gene, a *p1* paralog located ∼70 kb proximal to *p1* (Zhang et al., 2000). The *p1* and *p2* genes encode highly similar (95% identical) proteins (Zhang et al., 2000), and previous studies had shown that *p2/p1* chimeric genes are capable of producing pericarp pigment (Zhang et al., 2006; Wang et al., 2015). Therefore, we hypothesized that the red sectors observed on *p1-wwB54* ears represented activation of *p2* expression, possibly by CIs carrying and inserting a copy of the *p1* enhancer element in or near *p2*. The insertion of the *p1* enhancer would induce ectopic expression of *p2*, resulting in the red pericarp sectors observed on *p1-wwB54* ears. To test this hypothesis, we screened more than 1000 ears produced from plants carrying *p1-wwB54* and identified approximately 50 ears containing red sectors ranging in size from 1 kernel to whole ear (Figure 3). Classical genetic studies of *p1-vv* reversion show that the kernel pericarp (derived from the ovary wall) and egg cell share a cell lineage, and mutations that occur in pre-meiotic sporophytic cells can be transmitted through the female gametophyte (Wood and Brink, 1956). Kernels from independent red sectors were grown and propagated to establish a new allelic series of red pericarp types derived from the *p1-wwB54* allele.

**Figure 3.**
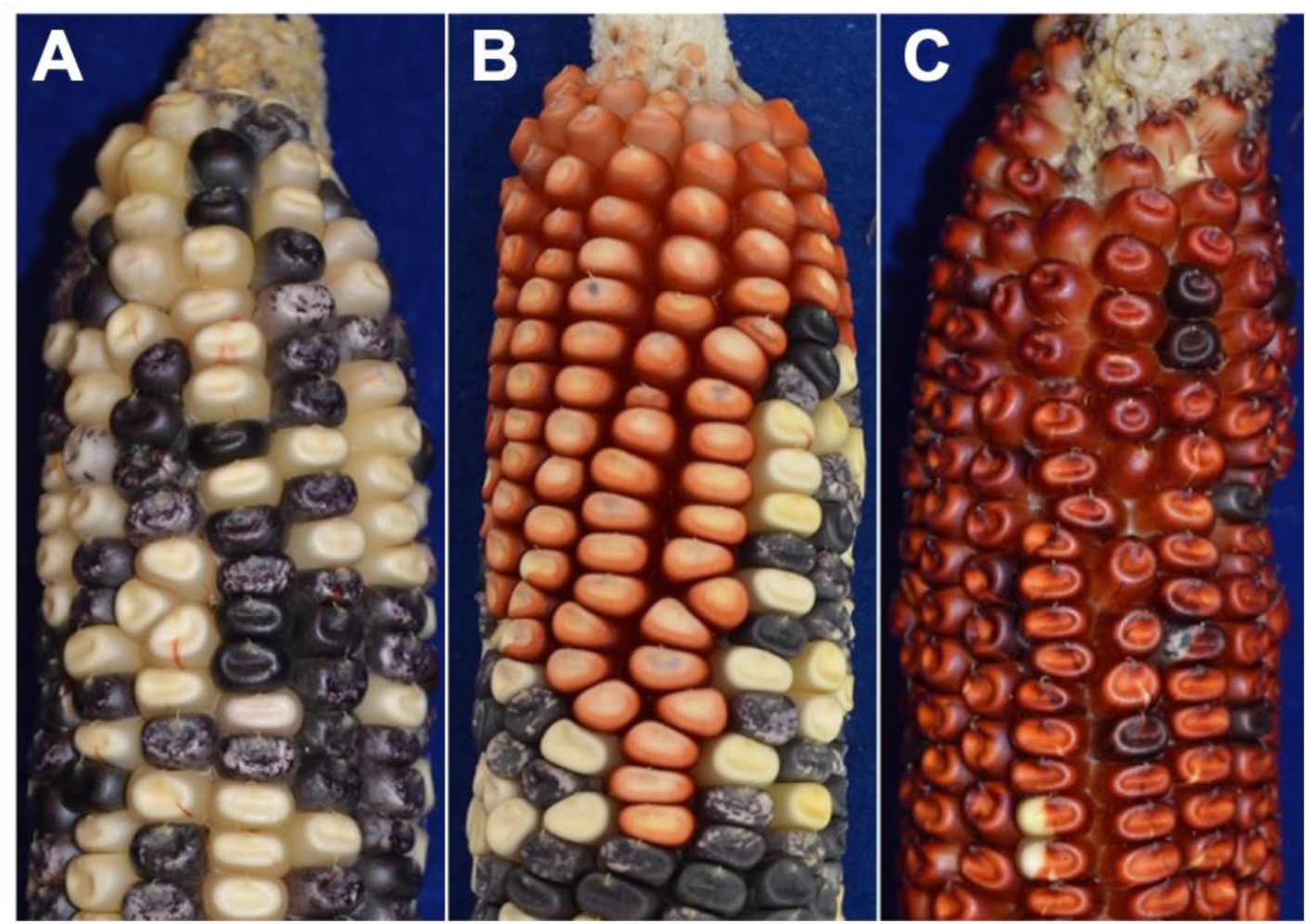
Screening for new CI alleles derived from *p1-wwB54*. A. Maize ear with typical *B54* phenotype with predominantly colorless pericarp, and small, infrequent red revertant sectors. B. Ear grown from *p1-wwB54* kernel, with a large multi-kernel red sector (upper) on an ear with otherwise typical *p1-wwB54* phenotype (lower portion of ear). C. Ear grown from *p1-wwB54* kernel with whole-ear red pericarp. Infrequent colorless sectors suggest ongoing instability of this novel allele, most likely due to *Ac* activity. In all ears, solid-colored and spotted kernels reflect *Ac-*induced excision of *Ds* element from *r1-m3::Ds* allele, resulting in sectors of purple kernel aleurone.

### Identification of Composite Insertions at *p2* locus

Using the screen described above, we isolated over 50 revertant alleles from *p1-wwB54*, and analyzed a representative sample in detail. First, using genomic PCR and Southern blot analysis (not shown), we determined that a large majority of revertant alleles tested do indeed carry new CIs inserted in or near the *p2* gene. For 24 cases, we mapped the sites of CI insertion by PCR using primers specific for *Ac* or *fAc* sequences, paired with primers in *p2* (Supplemental material 1). Reversed primers in *Ac* (*Ac-r*) were paired with reversed primers in *p2* (*p2-r*) to amplify the *Ac* junction, followed by a second PCR using *p2-f* plus *fAc-f* primers to amplify the *fAc* junction (Figure 4A and (Supplemental material 2). Figure 4B shows the PCR results from 10 CI alleles as examples. PCR products were sequenced and compared with *p2* genomic sequence to identify the precise insertion sites in 24 CI alleles: 10 cases contained CIs in the *p2* promoter region, while 14 cases had CIs in *p2* intron 2. Among these 24 CIs, 21 of them have the same orientation as shown in Figure 4A, with the *Ac* 5’ end closest to *p2* exon 3; 3 cases have the opposite orientation, in which the *fAc* 3’ end is closest to *p2* exon 3 (Figure 4C). By comparing the sequences of the *Ac* and *fAc* junctions in *p2*, we determined that each CI is flanked by an 8 bp Target Site Duplication (TSD) which is a characteristic feature of *Ac/Ds* insertion (Supplemental material 1). This finding confirms that the CIs are indeed generated by an *Ac/fAc* transposition event, consistent with the model proposed in Figure 2. Finally, we also identified 3 alleles in which the *Ac* element had excised from the CI, leaving behind a partial CI containing *fAc* and the *p1* sequences including the enhancer *f15*. This indicates that following CI formation, the *Ac* transposable element is still active and capable of subsequent independent transposition (Figure 4C and Supplemental material 1).

**Figure 4.**
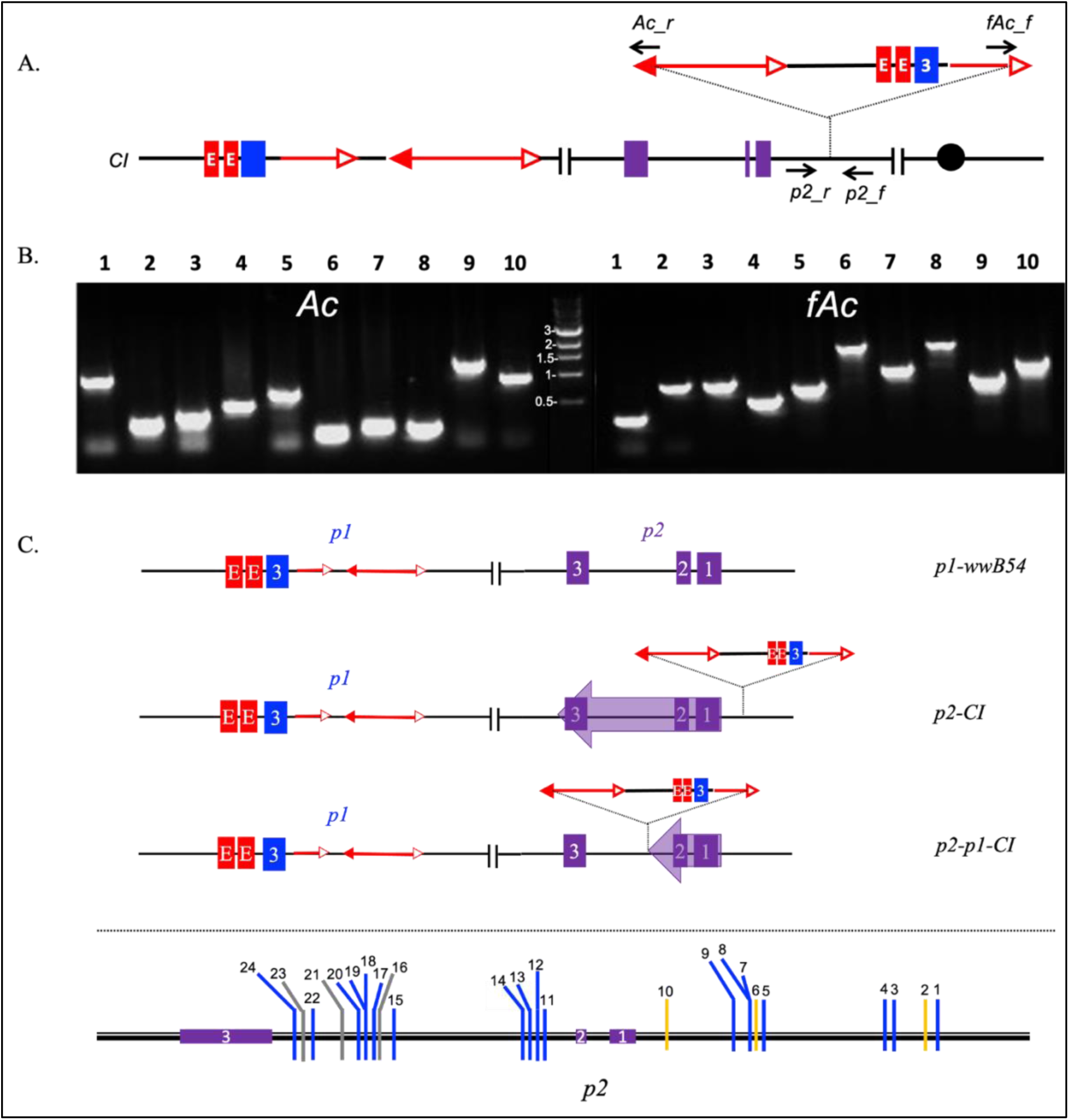
Identification of CI insertion sites. (A) The primer sets used for detection of CI target sites. *Ac_r* indicates a set of primers located on *Ac* 5’ end in a reversed orientation, *fAc_f* indicates primers located on *fAc* 3’ end in a forward orientation. *p2_r* and *p2_f* indicate primers in flanking *p2* sequence. (B) Results showing PCR amplification of *Ac* and *fAc* junction fragments from 10 independent CI alleles. Note that fragment sizes will vary depending on insertion site. Central lane is DNA size marker. (C) Map of CI insertions in *p2*. Upper panel: Diagrams of structures of progenitor *p1-wwB54*, and two types of *CI* alleles: *p2-CI* has CI insertion in *p2* promoter region, and *p2/p1-CI* has CI insertion into *p2* intron 2. Lower panel: Positions and orientations of 24 CIs in *p2*. In 10 cases, CIs are inserted in the *p2* promoter region (1-10), while 14 cases (11 – 24) have CIs inserted in *p2* intron 2. Blue lines indicate 18 insertions in the regular orientation shown in 4A; orange lines indicate 3 insertions in the opposite orientation; and 3 grey lines indicate 3 cases in regular orientation in which the *Ac* element has excised.

According to the model shown in Figure 2, DNA re-replication resulting from alternative transposition should generate CIs with varying sizes and sequence compositions. However, all CIs should contain sequences flanking the original *Ac* donor site, with *p1* 5’ sequences (upstream of *Ac)* fused to *p1* 3’ sequences (downstream of *fAc)* as shown in Figure 5A. Moreover, *p1* forward and *p1* reverse PCR primers which are divergent in *p1-wwB54*, should converge in each CI across the internal junction. To test this, we analyzed the internal structures of 10 independent CIs. The internal junction products were amplified by combinations of primers including *p1-r* + *p1-f* as shown in Figure 5A, and *Ac-f* + *p1-f* for those cases in which the internal junction was sufficiently close to the *Ac* 3’ end (Supplemental material 3). Due to the heterogeneity of CI length and structure, PCR was performed using series of *p1* forward and reverse primers to scan the region. In this way we isolated and sequenced the internal junctions of 10 independent CIs (Figure 5B and supplemental material 4); based on the internal junction sequences, we could surmise the structure of each case (Figure 5C). The 10 CIs range in size from 12.8 kb to 23.6 kb, including the *Ac* and *fAc* elements flanking each CI. In 6 out of the 10 alleles analyzed, the internal junctions contained microhomologies of 3 -19 bp, which are consistent with DSB repair via Non-Homologous End Joining (NHEJ) or Microhomology-Mediated End Joining (MMEJ) (Moore and Haber, 1996; McVey and Lee, 2008). The remaining 4 CI alleles contain additional filler DNA sequences inserted at each junction. These filler DNA sequences ranged in size from 4 to 50 bp and were apparently copied from nearby (within 100 bp) *p1* sequences, consistent with a template-switch mechanism as reported in previous studies (Wessler *et al*., 1990) (Supplemental Material 4).

**Figure 5.**
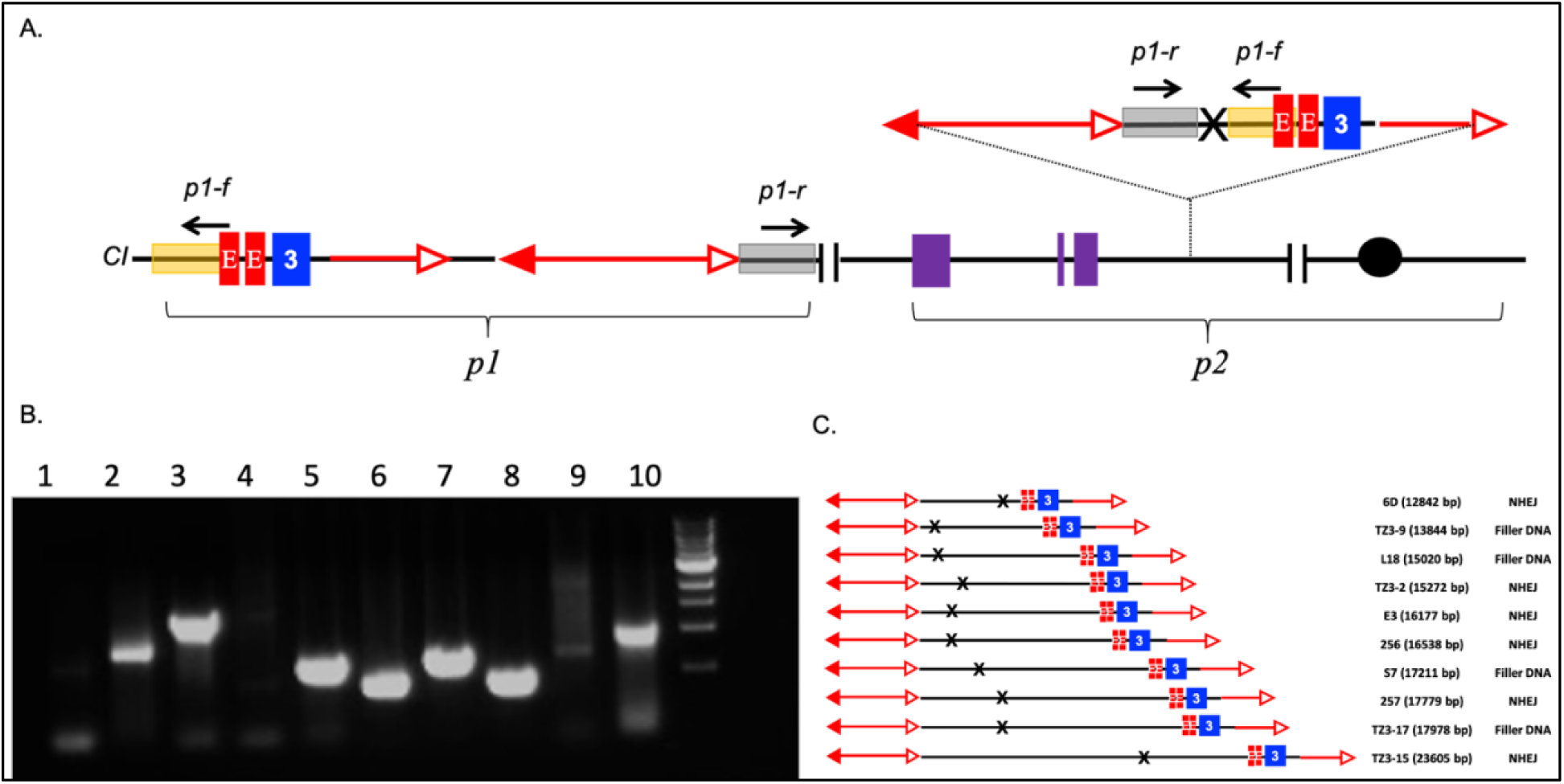
Identification of CI internal structures. A. Structure of representative chromosome containing the original *p1-wwB54* structure, and a new CI insertion into *p2* 5’ region. *p1-r* and *p1-f* represent sets of forward and reverse primers that are divergent in *p1-wwB54* (left), but convergent in CI (right). B. Results of PCR to amplify internal junctions of 10 CIs using *p1-f* and *p1-r* primers shown in 5A. The samples tested here correspond to the same 10 CI examples shown in figure 4B. Bands vary in intensity due to different PCR efficiencies using primers specific for each CI junction. C. CI structures in 10 representative alleles; the first column indicates CI names, with CI sizes in parenthesis; the second column indicates the DSB repair mechanism inferred from the junction sequences. CI insertions upstream of *p2* can induce transcription of the intact *p2* gene; while CI insertion into *p2* intron 2 can generate a chimeric *p2/p1* gene.

### Evidence that CI insertion drives *p2* expression

Importantly, all of the 10 CI cases examined contain 3’ *p1* sequences including transcriptional enhancer fragment 15 (indicated as red “E” box in Figure 5). This is consistent with the hypothesis that ectopic expression of *p2* in kernel pericarp in the CI-containing alleles is driven by the *p1* enhancer. A corollary to this hypothesis is that excision of the CI should result in loss of *p2* expression and reversion to the progenitor *p1-wwB54* phenotype. Excision of the CI as a macrotransposon may be expected, considering that it contains suitably oriented *Ac* and *fAc* transposons at each end. Indeed, many of the CI alleles exhibited variably-sized sectors of colorless and or less-pigmented pericarp (e.g. Figure 3C).

To test this hypothesis, we examined ears produced by *p2-S7*, an allele containing a 17.2 kb CI inserted upstream of *p2*. As shown in Figure 6, *p2-S7* conditions red kernel pericarp with some colorless sectors. Among ∼50 ears grown from *p2-S7* progenies, we identified one ear which had a large clonal sector of ∼20 kernels with near-colorless pericarp. Kernels from this sector gave rise to the stable mutant called *S7M*, which has a *p1-wwB54*-like phenotype (Figure 6A). We analyzed the structure of *S7M* by PCR using primers to amplify the original CI insertion site in *p2*, both *Ac* and *fAc* junctions with *p2*, and the internal CI junction (Figure 6B). The results (Figure 6B and 6C) show that in *S7M*, the CI excised from the target site as a macro-transposon, leaving behind the 8-bp TSD from the original insertion. These results show that *p2* expression was indeed a result of CI insertion, and removal of the CI eliminates the expression of *p2* and restores the phenotype of the progenitor *B54* allele.

**Figure 6.**
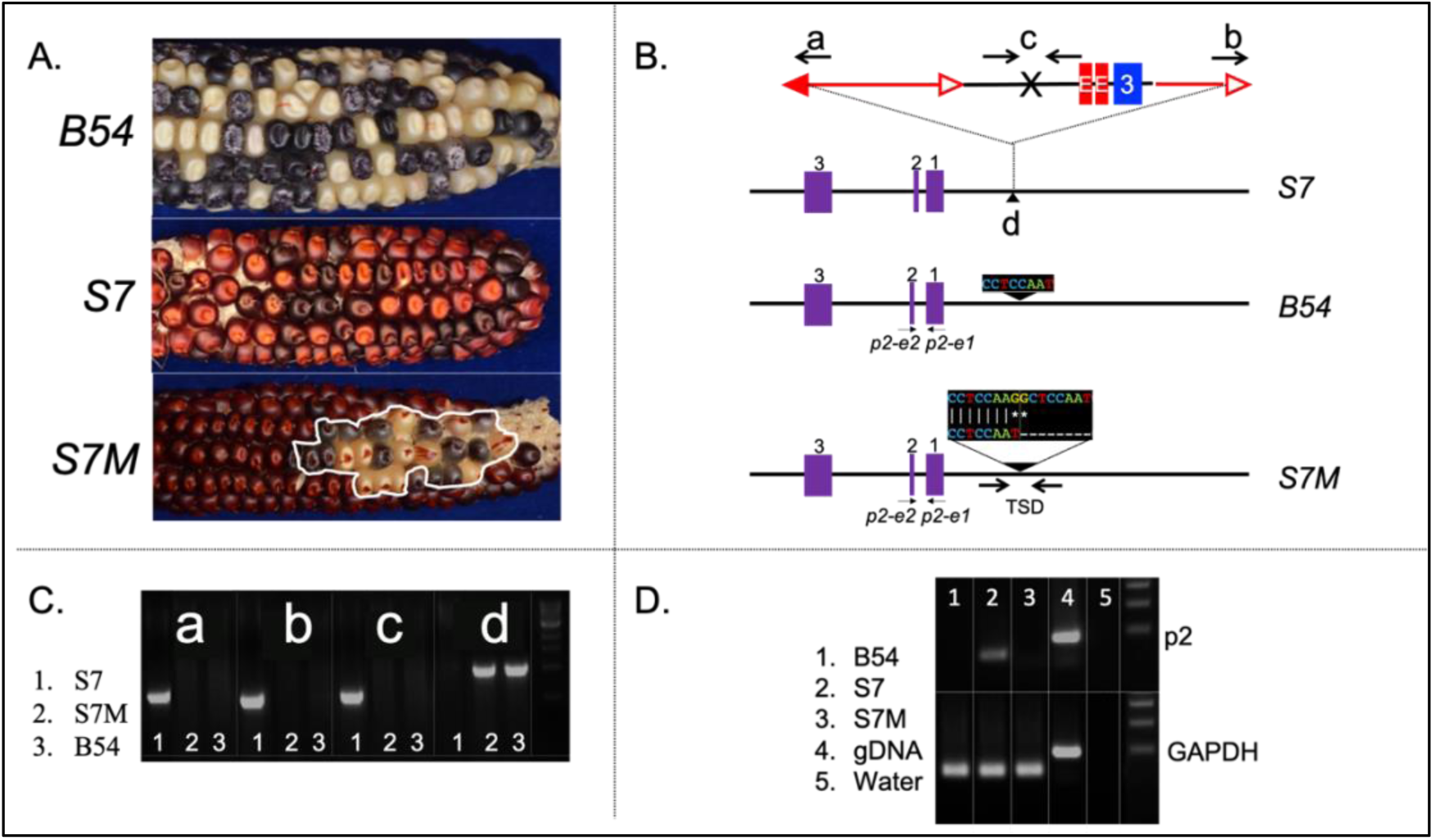
Isolation and analysis of CI-excision allele *S7M*. A. Phenotypes of *p1-wwB54, S7* and *S7M* (sector outlined in white). B. Structures of *S7, B54* and *S7M*. Letters and arrows indicate CI features analyzed by PCR; sequences indicate the target site in *B54*, and the TSD (Target Site Duplication) in S7M; arrows labeled p2-e1 and p2-e2 indicate the primers used in RT-PCR. C. PCR analysis of CI features in *S7, S7M*, and*B54*; a-d indicate corresponding features in S7 (Figure 6B). D. RT-PCR results showing the presence of *p2* transcripts in *S7*, and absence in *B54* and *S7M*.

To further test *p2* expression in the CI alleles, we measured *p2* transcript levels in *p1-wwB54, p2-S7* and *S7M* by RT-PCR (Figure 6D). Total RNA was prepared from developing kernel pericarp, reverse-transcribed into cDNA, and amplified with PCR primers located in *p2* exons 1 and 2 (*p2-e1* and *p2-e2* in Figure 6B and supplemental material 5). Primers complementary to the GAPDH gene were included as an internal control. The RT-PCR results showed that *p2* transcripts are detected only in the CI allele *S7*, and are undetectable in progenitor *p1-wwB54* and descendent *S7M* in which the CI had excised. PCR products were sequenced to confirm their origin from the *p2* gene (Supplemental material 6). These results confirm that the red pericarp phenotype was caused by the expression of *p2*, and that *p2* expression is dependent on presence of CI.

### P2-CI epiallele has altered DNA methylation

As noted above, some *p2-CI* alleles exhibited sectors and progeny ears with reduced pericarp pigment intensity. One case analyzed was derived from *p2-CI-E3* allele, which contains a 15.9 kb CI inserted in the 5’ region of *p2*. This variant (termed *E3M*) was isolated from a single kernel in a small sector of light orange pericarp on an otherwise red *E3* ear (Figure 7A). Progeny plants grown from this kernel have distinctly lighter orange kernel pericarp, indicating a heritable reduction in *p2* expression in *E3M*. However, unlike CI-excision allele *S7M*, PCR analysis of the *Ac, fAc* and internal junctions showed that *E3M* does not have any structural variations in the CI target site (Figure 7B and 7C). We hypothesized that the *E3M* dilute-pigment phenotype was caused by epigenetic change(s) rather than structural variation. Epigenetic variations such as DNA methylation are known to be correlated to changes in gene expression (Assaad *et al*.,1993). Therefore, we conducted Bisulfite Sequencing (BS) of seven targeted regions in *p1* and *p2* to analyze DNA methylation at single-base resolution. We first examined methylation of the *f15* enhancer fragment. Because both the background *p1* gene and *CI-E3* each contain two tandem copies of *f15*, we used primers specific to the second copy downstream of *p1;* in *CI-E3*, this copy is located nearest the internal junction and nearest the *p2* transcription start site (Figure 7D, box 1). To improve product quality and yield, initial products were re-amplified using internal nested primers (Supplemental Material 7). PCR products were sequenced to identify methylated cytosines in a 100 bp segment at the end of the enhancer in both background *p1* and the CI. The results showed that all 24 cytosine residues in this region are unmethylated in *B54, E3* and *E3M*. (Figure 7E and Supplemental material 8).

**Figure 7.**
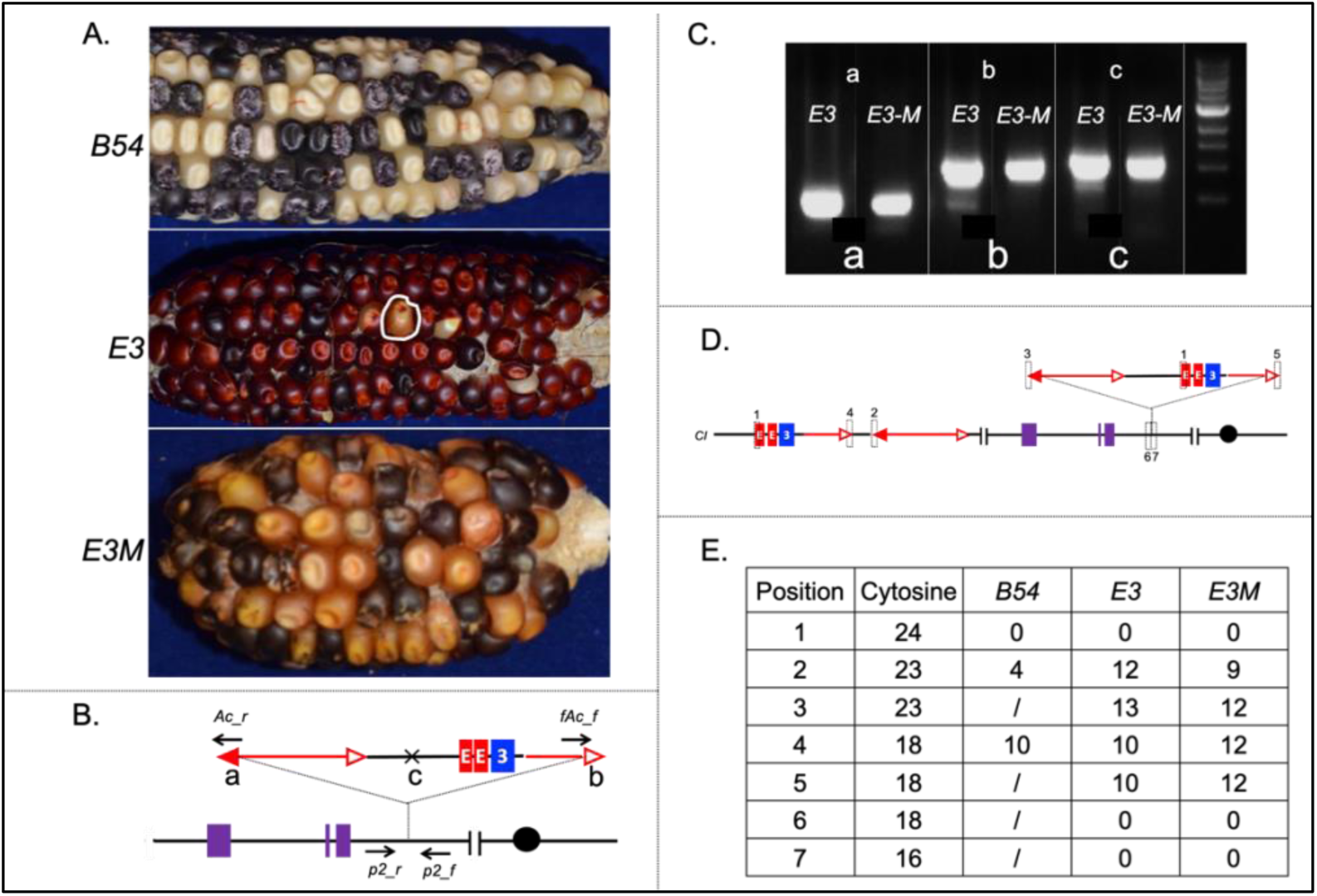
Epiallele *E3M* derived from CI *E3*. A. Ear and kernel pericarp phenotypes of *B54, E3* and *E3M*. Epiallele *E3M* originated from pale kernel outlined in *E3*. B. Structure of *CI-E3* allele showing locations of PCR primers used to confirm *E3* and *E3M* CI insertions. C. PCR results of *Ac, fAc* and the internal junctions corresponding to 7B (a-c). D. Genomic segments targeted for bisulfite sequencing to analyze cytosine methylation. Dotted boxes numbered 1 – 7 indicate sites analyzed. E. Summary of cytosine methylation data in the regions annotated in panel D. The first column indicates the genomic positions corresponding to 7D, the second column indicates the total number of cytosines in each segment; remaining columns indicate the number of methylated cytosines in *B54, E3* and *E3M*. “/” indicates absence of that sequence in the *B54* allele.

We also analyzed cytosine methylation of the first 100 bp of both 5’ *Ac* and 3’ *fAc* termini in CI, and compared with the same regions in the *Ac* and *fAc* elements located in the linked *p1-wwB54* locus (hereafter referred to as background *Ac* and *fAc)*. We compared the results among alleles based upon their lineage relationship: the *p1-wwB54* allele is progenitor of *CI-E3*, which itself gave rise to *CI-E3M*. In the first 100 bp 5’ *Ac*, there are 23 cytosines. In the background *Ac* (Figure 7D, box 2), *B54* has 4 methylated and 19 unmethylated cytosines in this region, while *E3* and *E3M* have 12 and 9 methylated cytosines, respectively. In the CI *Ac* (3 in Figure 7D), *E3* has a net +1 additional methylated cytosine compared to the background *Ac*; this results from 1 de-methylation and 2 *de novo* methylations. In *E3M*, the CI *Ac* has 3 *de novo* methylated cytosines compared to the background *Ac* (Figure 7E and supplemental material 8). Next we examined the first 100 bp of *fAc* which has 18 cytosine residues; in the background *fAc* (Figure 7D, box 4), *B54* has 10 cytosines methylated, while *E3* and *E3M* have 10 and 12 methylated residues, respectively, in the corresponding region. In the CI *fAc* (Figure 7D, box 5), *E3* has the same methylation pattern as the background, while *E3M* has 1 de-methylated cytosine and 1 *de novo* methylation (Figure 7E and Supplemental material 8).

Finally, we analyzed the 50 bp *p2* sequences flanking both *Ac-CI* (Figure 7D, box 6), and *fAc-CI* (Figure 7D, box 7); the result showed that the 18 cytosines in the *Ac-CI* flanking region and the 16 cytosines in the *fAc-CI* flanking region are unmethylated. Together, these results indicate that changes in cytosine methylation associated with the *CI-E3M* epiallele were detected only in the *Ac* and *fAc* sequences, and did not appear to spread into the *f15* enhancer, nor outward into p2 flanking sequence.

These results showed that methylation does not change dramatically between the E3 and E3M alleles in the segments analyzed. However, a recent report indicates that changes in methylation at a single CpG can influence transcription factor binding (Yang et al, 2020). Therefore, although we cannot rule out that the observed differences in methylation of *E3* and *E3M* are not associated with differential expression of *p2*, it is possible that the specific methylations observed may impact *p2* expression.

## Discussion

Transposable elements comprise a large proportion of most eukaryotic genomes, and are usually considered to be selfish elements providing little or no benefit to the host genome (Orgel et el, 1980). Many studies have shown that TEs often have deleterious effects such as disrupting gene structures and modifying the epigenetic features near their insertion sites (Hollister and Gaut, 2009). However, recent studies have shown that TEs can modify coding sequences and regulate gene expression to potentially increase the fitness of the host (Chuong et al., 2017). Specifically, transposable elements can perform enhancer-like activities in eukaryotic genomes. For example, in the human genome, widespread enhancers are overlapping with TEs (Cao et al., 2019), and experimental data validated that a subset of TE-enhancers played important roles in gene regulation in early mouse development (Todd et al., 2019). These studies supported the long-standing hypothesis that TE domestication is important in eukaryotic genome evolution and that some TEs play integral roles in regulating gene expression. However, most of these studies were focused on Class I TEs in animal and human systems (Sundaram and Wysocka, 2020). In this study, we identified a new mechanism by whichClass II TEs can influence gene regulation in maize. We demonstrate that, in addition to evolving into regulatory elements over time, TEs can induce sudden changes in gene expression by changing the copy number and configuration of existing enhancer elements.

In previous studies, we have described the mechanism of RET-induced DNA re-replication in maize (Zhang et al., 2014). This re-replication process is initiated by *Ac/fAc* transposition which generates a rolling circle replicon to replicate the TE and flanking sequences an additional round in the same cell cycle. Here we show that a regulatory element, enhancer *f15*, can be duplicated and transposed in this process. We screened ears produced from *p1-wwB54* and isolated approximately 50 ears containing red sectors ranging in size from 1 kernel to the whole ear. We mapped the insertion sites for 24 of them using primers located in the *p2* promoter region and intron 2 sequence. Ten CIs were inserted in the upstream sequences of *p2*, while 14 insert in *p2* intron 2. A few insertion hotspots were observed; for example, we detected 5 CI insertions in a <200 bp region upstream of *p2* (positions -3188 to -3364). Similarly, 4 CIs inserted into a <100 bp region in *p2* intron 2 (positions 8017-8102). Moreover, some CIs insert very close to each other, or even at the same site (L12 and S7; TZ3-1 and TZ3-12; S10 and TZ3-17). We did not detect any clear sequence signatures in these hotspots (Vollbrecht et al., 2010) (data not shown); possibly insertion site preference may be influenced by epigenetic modifications. We analyzed the detailed structures of 10 CIs ranging in size from 12.8 kb to 23.6 kb. All were composed of *Ac* and *fAc* elements enclosing duplications of sequences flanking the donor elements. These duplications were joined together at internal junctions with sequences characteristic of fusion by NHEJ, accompanied by presence of filler DNA sequences in half the cases. These structures are all consistent with a model of CI formation by DNA re-replication induced by RET (Zhang et al., 2014)

Activation of *p2* expression by the enhancer-containing CI was confirmed by analysis of a particular case, *S7M*, in which the complete CI excised as a macro-transposon. CI excision resulted in heritable loss of kernel pericarp pigmentation and elimination of *p2* RNA, proving that the red pericarp phenotype was caused by CI-induced *p2* expression. Another variant allele (*E3M)* which specified orange pericarp phenotype was analyzed and found to have some DNA methylation changes in the terminal sequences of the CI *Ac* and *fAc* elements. It is not clear whether these methylation differences are responsible for the differences in *p2* expression; however, a previous study showed that TE DNA methylation can impact the expression of nearby genes (Wittmeyer et al., 2018).

These results indicated that alternative transpositions are not only able to increase the number of regulatory elements, but also modify their distance to the target genes, and thereby regulate gene expression patterns. This mechanism is meaningful in plant development and genome evolution. In our case, depending on the length of the re-replication, the CIs enlarged the genome by various sizes. Furthermore, the target *p1/p2* locus plays a central role in regulating phlobaphene biosynthesis in maize tissues (Grotewold et al, 1994). A recent study has demonstrated a direct correlation between phlobaphene accumulation and kernel pericarp thickness, as well as reductions in mycotoxin contamination on maize kernels. Therefore, the ectopic expression of *p2* induced by CI alleles could be beneficial to the plants (Landoni et al., 2020).

Although in the evolutionary process TEs proliferate in the genomes and contribute to a large portion of repetitive sequences, most of the TEs are epigenetically silenced. Evidence showed that TEs sequences are heavily methylated in both plant and animal genomes (Quadrana and Colot, 2016; Hollister et al. 2009; Aravin el al., 2008). Moreover, the silencing signal can spread beyond the TE and affect the flanking sequence and nearby genes (Noshay et al., 2019). These silenced TEs are immobile or reduced in transposition potential, and thus are hardly able to generate large genome rearrangements. In maize, the *Mu* element and the *Ac/Ds* elements have been characterized as active TE families that tend to land in low methylation regions with open chromatin structures (Springer et al, 2018). In this study, we show that after CIs inserted to the target sites, the *Ac* element is still active, it can excise and insert to other loci as standard transposition. Furthermore, the mutant from *S7* allele indicates that the CI can move as a macrotransposon. Although we did not detect transposition of the CI to other loci in *S7M*, it is quite possible that a macro-transposon of this size can excise and insert in the genome. Moreover, mutation or loss of the *CI Ac* 3’ end would prevent independent excision of the *Ac* element, converting the complex *CI* macro-transposon into a single mobile element. This provides one plausible mechanism for sequence acquisition by TIR elements. For CI’s that contain functional enhancers as described here, such cases may be considered as authentic *Controlling Elements* as originally described by McClintock (McClintock, 1956) In this study, we used *Ac/fAc* and the *p1/p2* loci as examples to reveal the potential regulatory role of alternative transpositions in plant development. It should be noted that because of the system, our screening of the CI insertions was biased toward the visible marker of red phenotypes. In fact, CIs can insert into any locations in the genome, not necessarily producing an observable phenotype. Moreover, because there are thousands of TIR TEs in the maize genome (Su et al, 2019), it is possible that similar RET events occur at other loci involving other TE systems and and affecting other genes. Therefore, a much more significant impact of alternative transposition on gene expression can be postulated. In addition to maize, alternative transpositions have also been identified in snapdragon *Tam3* elements (Martin and Lister, 1989) and *Drosophila P* elements (Gray et al., 1996). This mechanism could potentially be an important source of modifications of regulatory networks in both plant and animal genomes. In conclusion, our study demonstrated a new mechanism for gene regulation by transposable elements.

## Acknowledgments

We thank Terry Olson for technical assistance, and Drs. Jianbo Zhang and Dafang Wang for suggestions on the experiments. This research is supported by the USDA National Institute of Food and Agriculture Hatch project number IOW05282, and by State of Iowa funds.

## Author contributions

W.S., T.Z., and T.P. conceived and designed the experiment; W.S. and T.Z. performed the experiments; W.S. and T.P. wrote the paper.

---

**Supplemental material 1.**
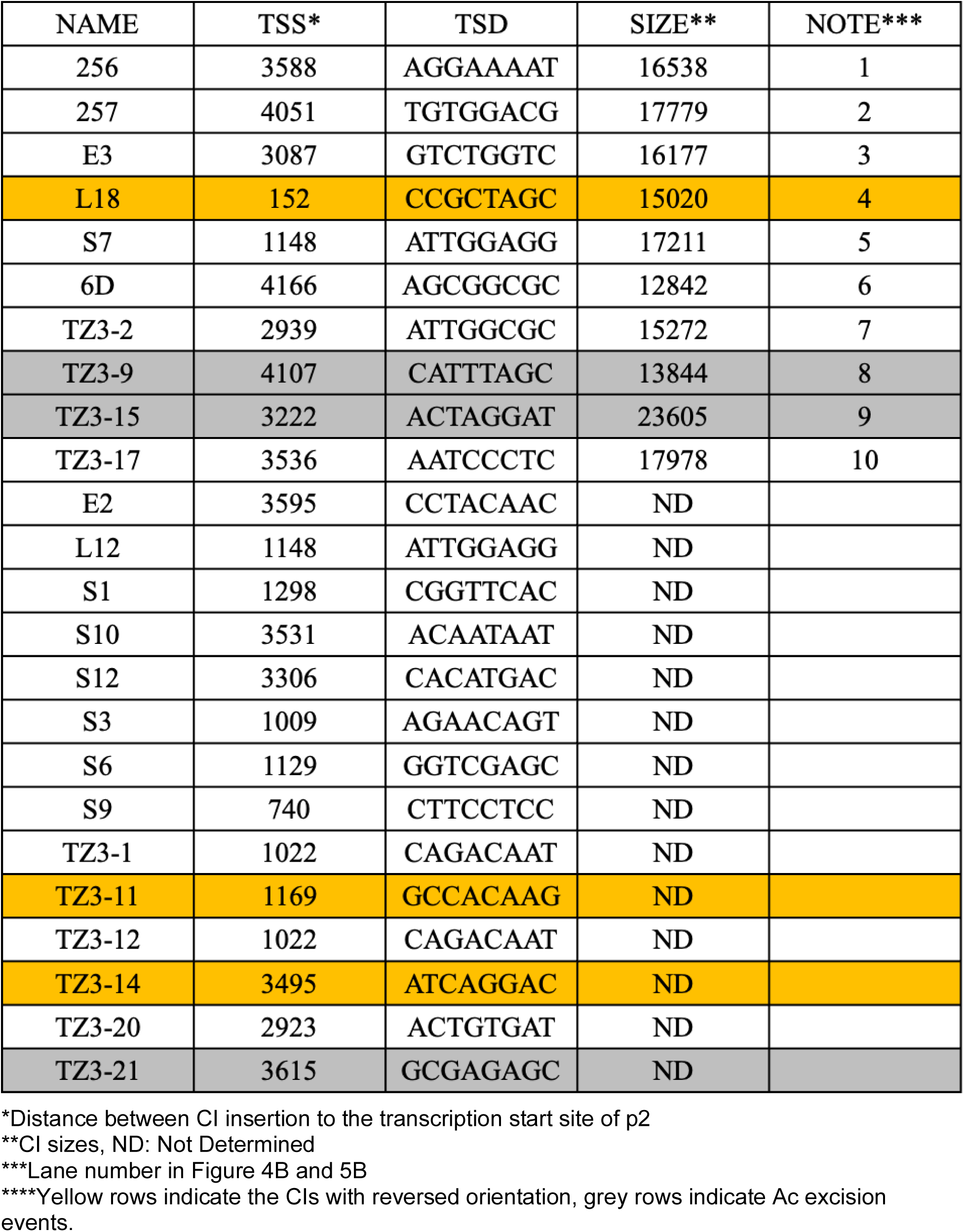
Map of 24 alleles

**Supplemental material 2.**
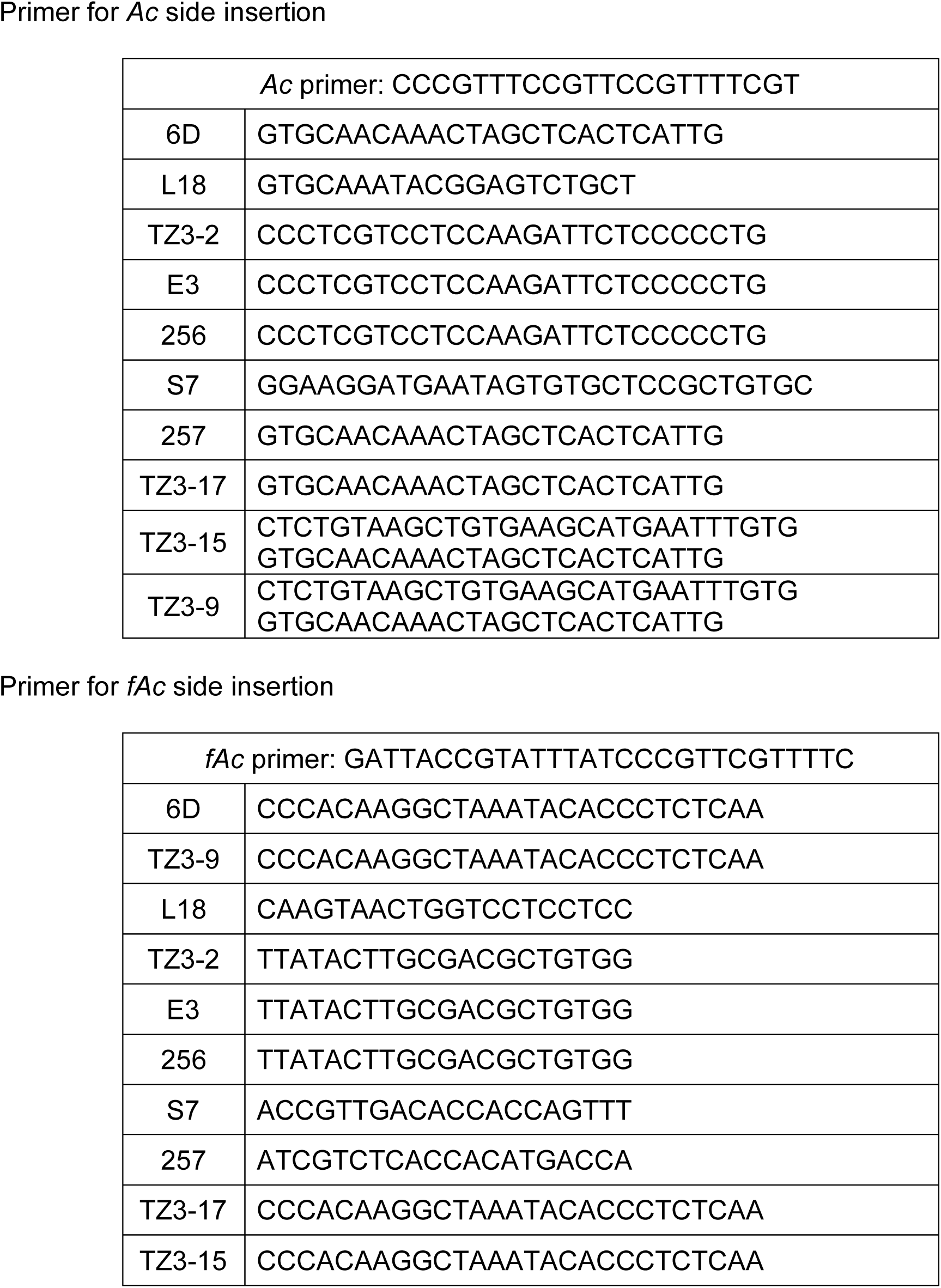
Primers for detecting *Ac* and *fAc* junctions.

**Supplementary material 3.**
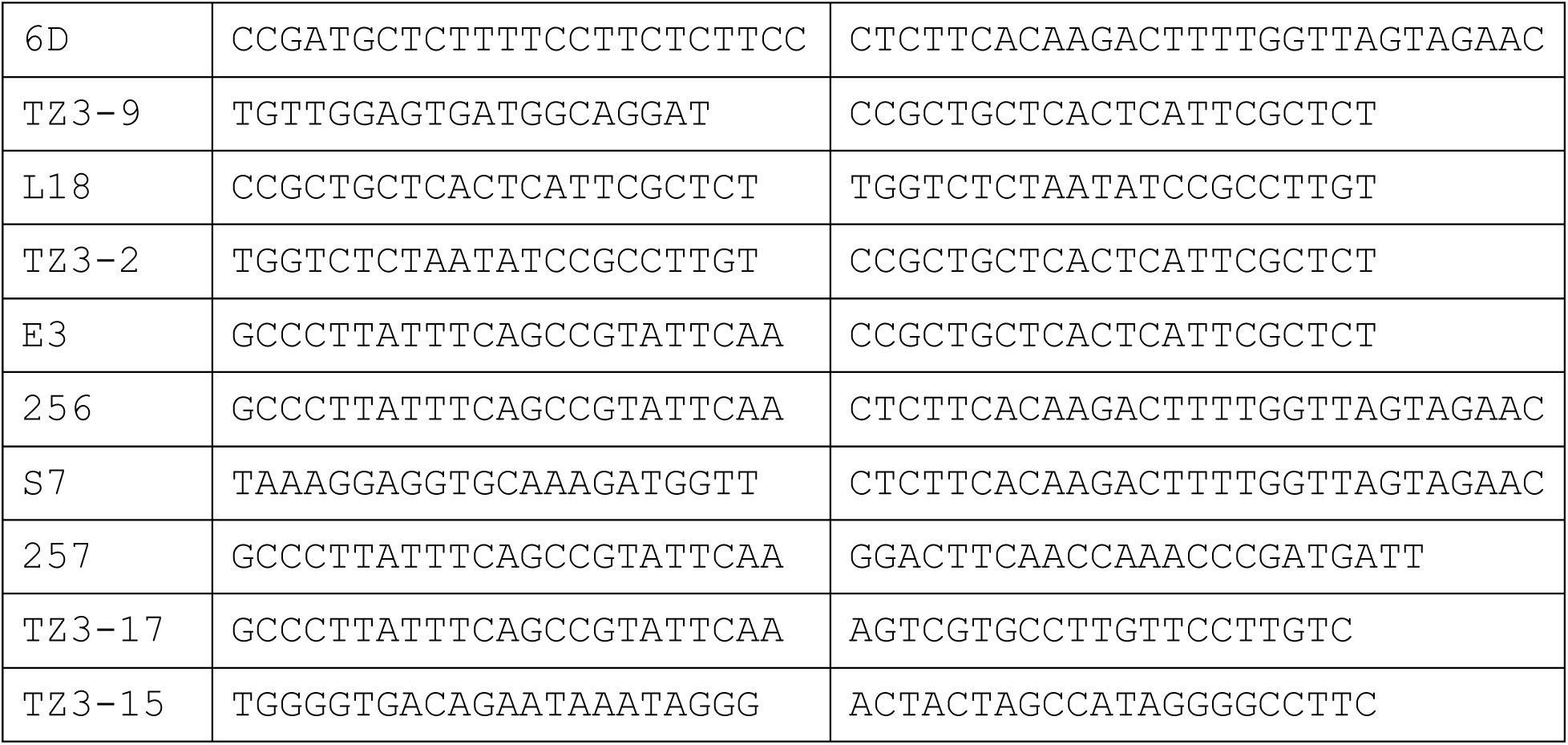
Primers for internal junctions

**Supplementary material 4.**
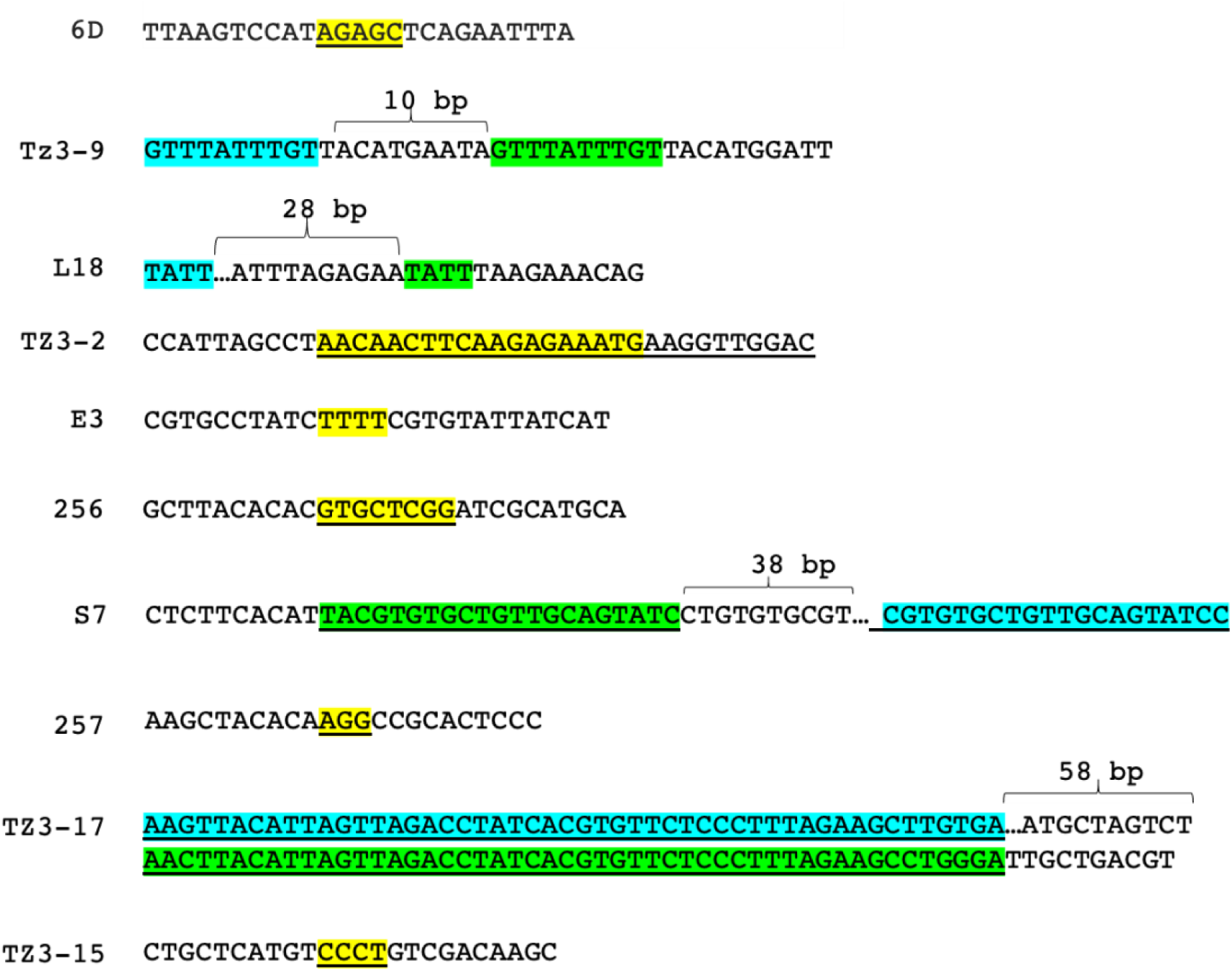
Junction sequence of the ten alleles Microhomology sequences (yellow), Filler DNA (green) and the original *p1* sequences that produced the filler DNA (blue).

**Supplementary material 5.**
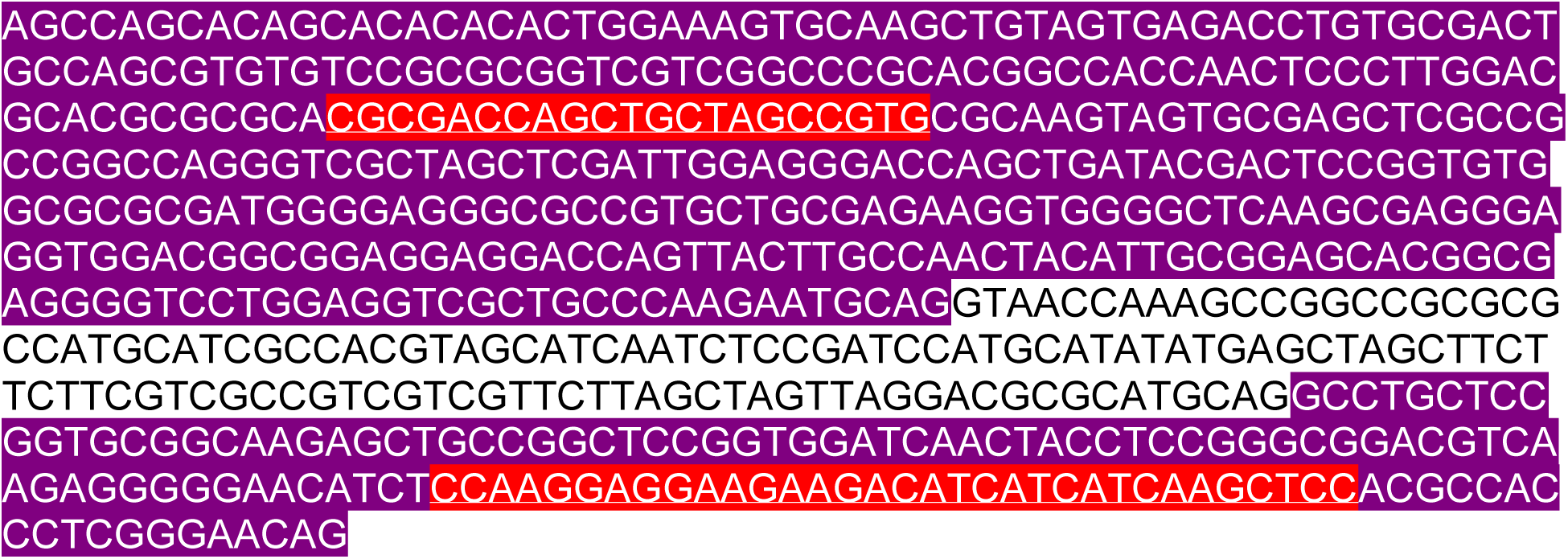
Primers for RT-PCR: Purple sequences indicate the exon 1 and exon 2, red underlined sequences indicate the primers used for RT-PCR, and the white sequence in the middle indicates the intron 1 of *p2*.

**Supplementary material 6.**
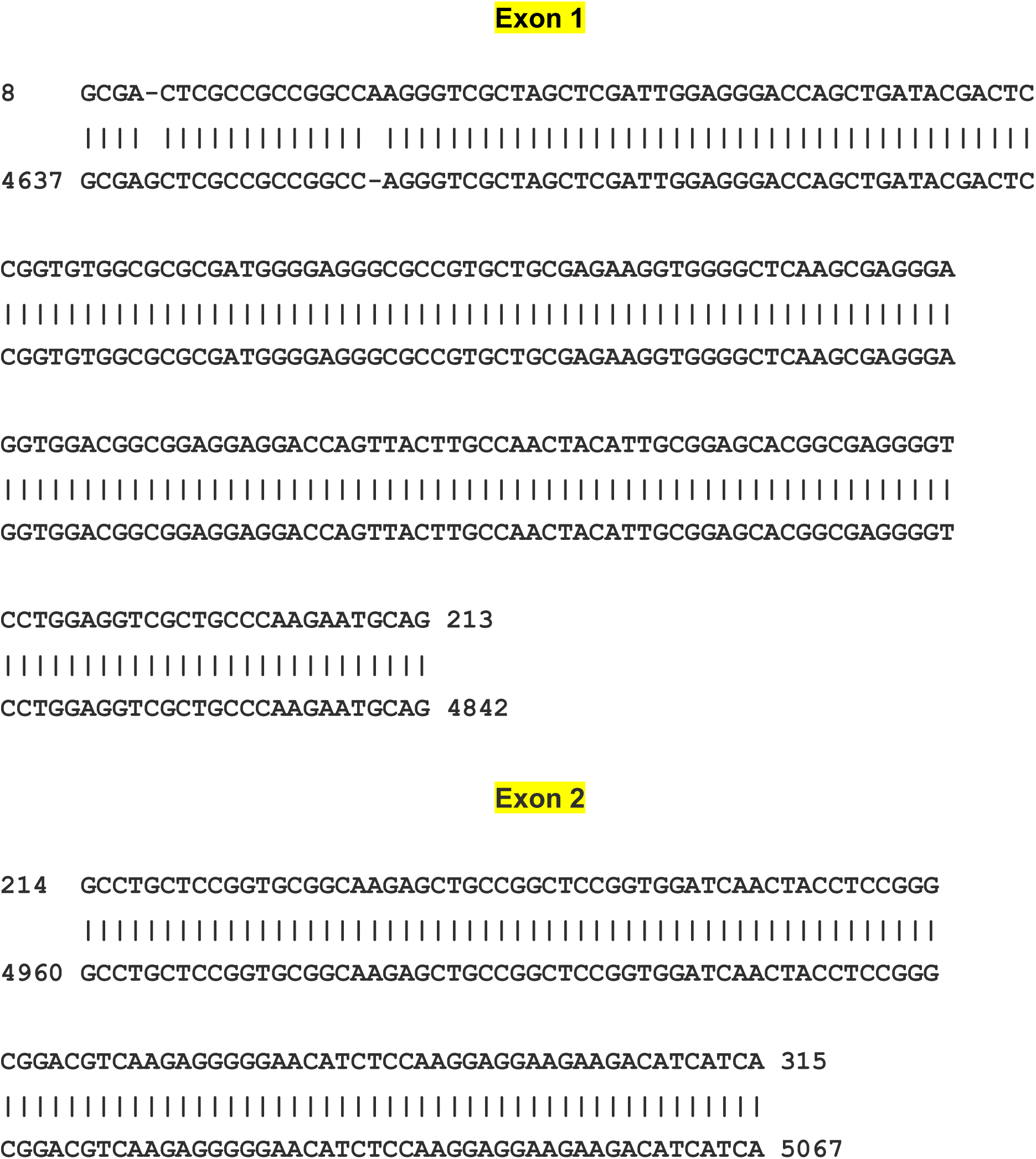
RT-PCR sequence aligned to *p2* exon 1 and exon 2, the upper line indicates RT-PCR product from *S7*, and the lower line indicates the *p2* sequence.

**Supplementary material 7.**
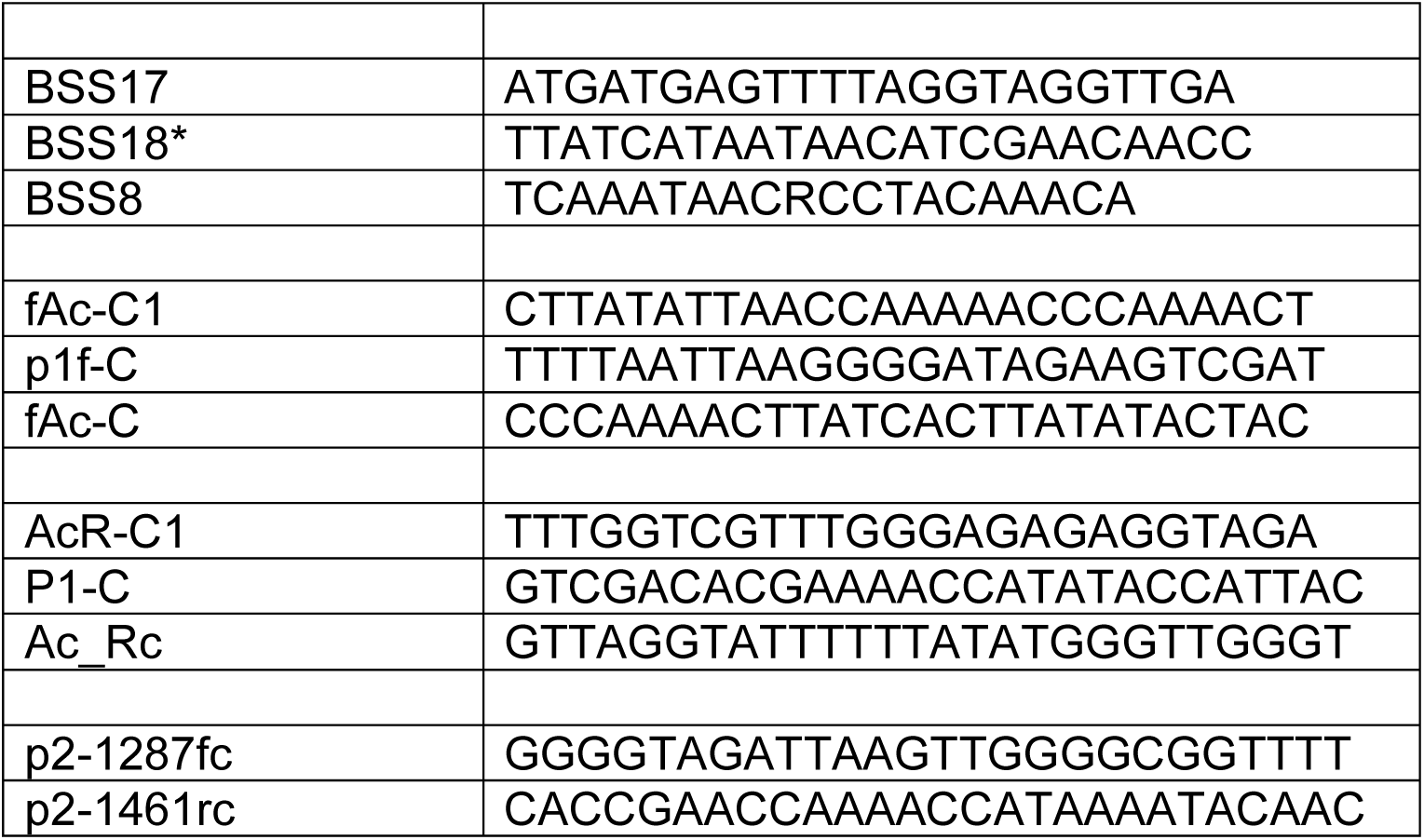
Primers for Bisulfite Sequencing

**Supplementary material 8.**
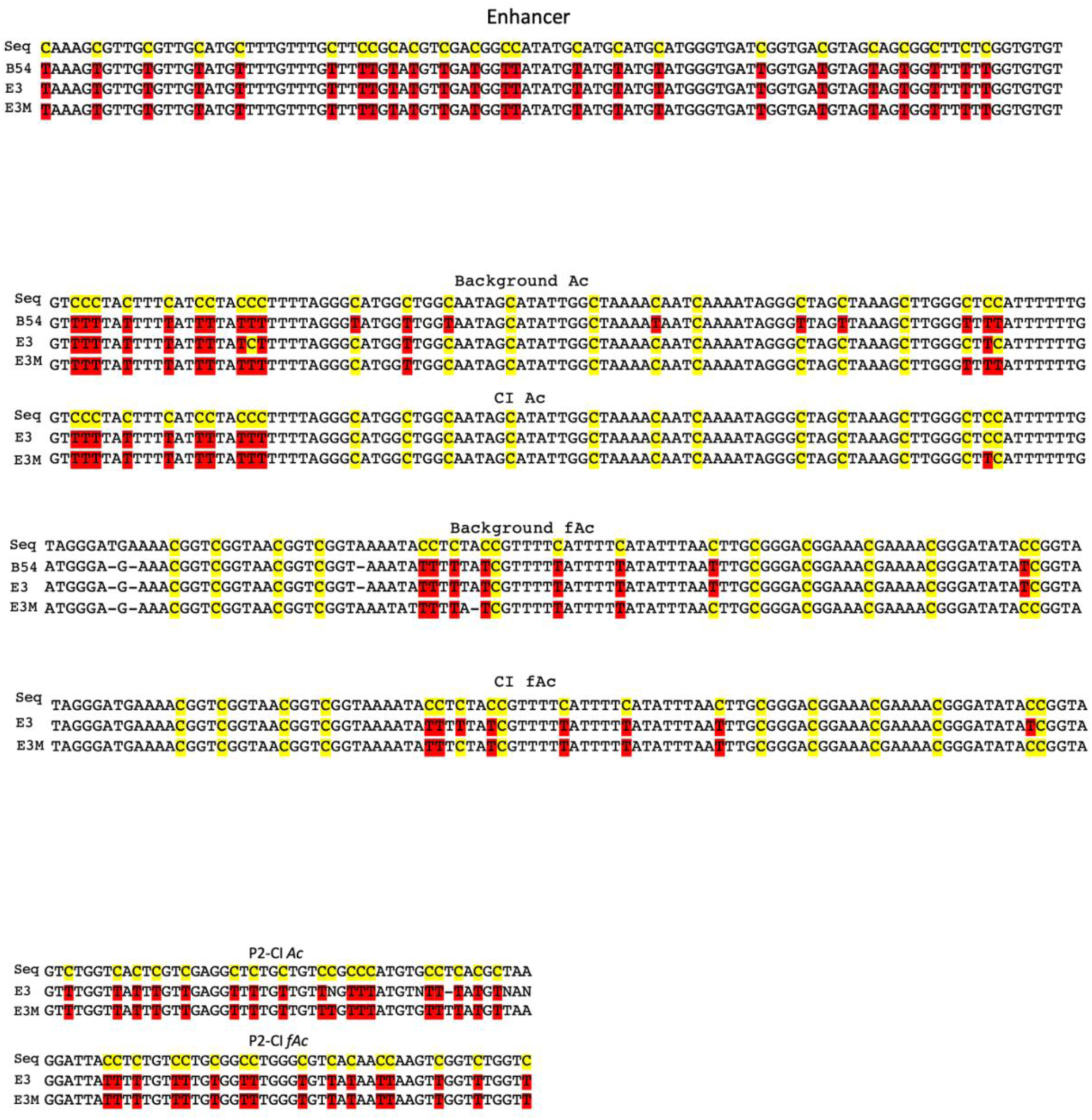
Bisulfite sequencing alignments: Cytosines in the sequence are labeled in yellow, unmethylated cytosines (T) are labeled in red and methylated cytosines remain in yellow.

